# Polymer Simulations of Heteromorphic Chromatin Predict the 3-D Folding of Complex Genomic Loci

**DOI:** 10.1101/380196

**Authors:** Adam Buckle, Chris A Brackley, Shelagh Boyle, Davide Marenduzzo, Nick Gilbert

## Abstract

Chromatin folded into 3-D macromolecular structures is often analysed by 3C and FISH techniques, but frequently provide contradictory results. Instead, chromatin can be modelled as a simple polymer comprised of a connected chain of units. By embedding data for epigenetic marks (H3K27ac), genomic disruptions (ATAC-seq) and structural anchors (CTCF) we developed a highly predictive heteromorphic polymer (HiP-HoP) model, where the chromatin fibre varied along its length; combined with diffusing protein bridges and loop extrusion this model predicted the 3-D organisation of genomic loci at a population and single cell level. The model was validated at several gene loci, including the complex *Pax6* gene, and was able to determine locus conformations across cell types with varying levels of transcriptional activity and explain different mechanisms of enhancer use. Minimal *a priori* knowledge of epigenetic marks is sufficient to recapitulate complex genomic loci in 3-D and enable predictions of chromatin folding paths.

## Introduction

Chromatin fibre folding in cells is dictated by a vast number of interactions between nucleosomes, chromatin binding proteins, and structural components such as CTCF/cohesin loops, as well as the inherent structure of the underlying fibre. Chromatin is far from a bland homomorphic fibre, rather it is a structurally heterogeneous material which is frequently disrupted at transcriptional hotspots^1,2^, and is thought to be locally compact in inactive regions. The ENCODE project^3^ comprehensively mapped the distribution of epigenetic and structural features in different human and mouse cell lines. Many of these marks are surrogates for transcriptional activity that can impact on local chromatin fibre structure. Previously we developed a polymer modelling scheme based on the assumption that chromosome organisation is driven by the formation of bridges by multivalent protein complexes (the transcription factor, or TF model^4–6^). For example, complexes of TFs and polymerase form enhancer-promoter interactions to organise active regions, while PRC (polycomb repressor complex) or HP1 proteins might arrange inactive and repressed regions. This model can predict large-scale organisation such as chromatin domains and compartments^6^, and has been successful in describing some genomic loci at higher resolution (e.g. the α and β-globin loci^5^). We also recently combined the TF model with the popular loop extrusion (LE) model for chromosome organisation^7^, which explains features of chromatin loops mediated by cohesin and CTCF^8,9^. While this strategy successfully predicts large-scale features of genome organisation, we show below that it cannot accurately predict the folding of the complex *Pax6* genomic locus at high resolution, which we probed experimentally at different levels using fluorescence *in situ* hybridization (FISH) imaging and Capture-C. *Pax6* is surrounded by constitutively expressed genes and multiple enhancers, providing a paradigm for complex genetic interactions^10–12^.

Our previous models use a simple bead-and-spring polymer to represent the chromatin fibre, and as such assume that this has a uniform structure. We speculated that certain histone modifications would be indicative of disrupted chromatin with decreased linear fibre compaction; to this end we developed a new predictive heteromorphic polymer (HiP-HoP) model. Simulations of these “heteromorphic” chromatin fibres gave a much better recapitulation of locus conformation at transcriptionally active regions of the genome, providing a universal model for chromatin fibre folding which could potentially be applied to map 3-D structures genome-wide in the future — as examples here we studied the *Pax6*, globin and *Sox2* loci. Unlike other widely used “inverse modelling” approaches for predicting 3-D chromatin folding, such as the recent polymer-physics-based approach PRISMR^13^, or previous approaches based on Markov Chain Monte Carlo or constrained molecular dynamics^14–18^, the present scheme does not rely on *any* fitting to pre-existing 5C or HiC data, making it applicable to a wider variety of experimental situations, for instance to investigate the 3-D conformation of rare or hard-to-obtain tissues and cell types.

## Results

### Activity states of *Pax6* show differential epigenetic marks and CTCF binding

In the present work we set out to develop a universal approach for modelling chromatin fibre folding with limited experimental knowledge, based only on extensive freely available data generated from the ENCODE project. To develop this strategy we investigated the folding of 5 Mb around the *Pax6* locus using three different immortalized cell lines that expressed *Pax6* at different levels (Fig. S1), referred to as Pax6-OFF, ON and HIGH cell lines. *Pax6* is flanked by two constitutively expressed housekeeping genes, with enhancer elements within the *Pax6* gene itself, and at regions ~50 kb upstream and ~95 kb downstream — these are referred to below as the up and downstream regulatory regions (URR and DRR respectively^11,19^). The histone modification H3K27ac (Fig. S1), usually associated with enhancers and transcriptional activation, was found at the gene and distal enhancers when *Pax6* was active, and these regions at enhancers broadened significantly in HIGH activity cells. Surprisingly, despite large differences in *Pax6* transcription, CTCF and Rad21 binding across the locus did not vary significantly between the cell lines (Fig. S1), but additional CTCF bound in close proximity to the *Pax6* promoters in *Pax6* expressing cells.

### Active epigenetic marks predict locus folding only in some cell lines

Our previous modelling work^5–7^ gave good predictions of larger-scale (domain and compartment level) chromosome organisation. To test whether this scheme can also predict folding of complex genetic loci at higher resolution, we performed simulations for the *Pax6* locus. This model, where a chromosome region is represented as a bead-and-spring polymer (with each bead representing 1 kb), combines two views on what drives chromatin conformation (see Fig. 1a for a schematic). First, the TF model^5,6^ postulates that promoter-enhancer interactions are mediated by diffusing protein complexes which form molecular bridges between their binding sites. Here we began by assuming that TFs bind H3K27ac regions, and switch back and forth between a binding and a non-binding state. Switching models post-translational modifications, active protein degradation, or programmed polymerase unbinding after transcription termination^7,20^; it enables simultaneously strong TF binding and fast turnover of bound TFs (as observed by photobleaching experiments^21^), and drives the system away from equilibrium. Second, the LE model^8,9^ views cohesin and CTCF as the structural organisers of the genome, with cohesin forming chromatin loops via an extrusion mechanism that could be transcription dependent^22^. LEs stop if they encounter a CTCF site which has a binding motif oriented towards the direction of extrusion; this enables stable looping between CTCF sites with binding motifs which are in a convergent, but not divergent arrangement^23^. We used the model to generate an ensemble of locus conformations representing a population of cells and from this extracted both 3C-like information, and single cell simulated FISH data (see Methods for details of combined TF+LE simulation scheme).

**Figure 1.**
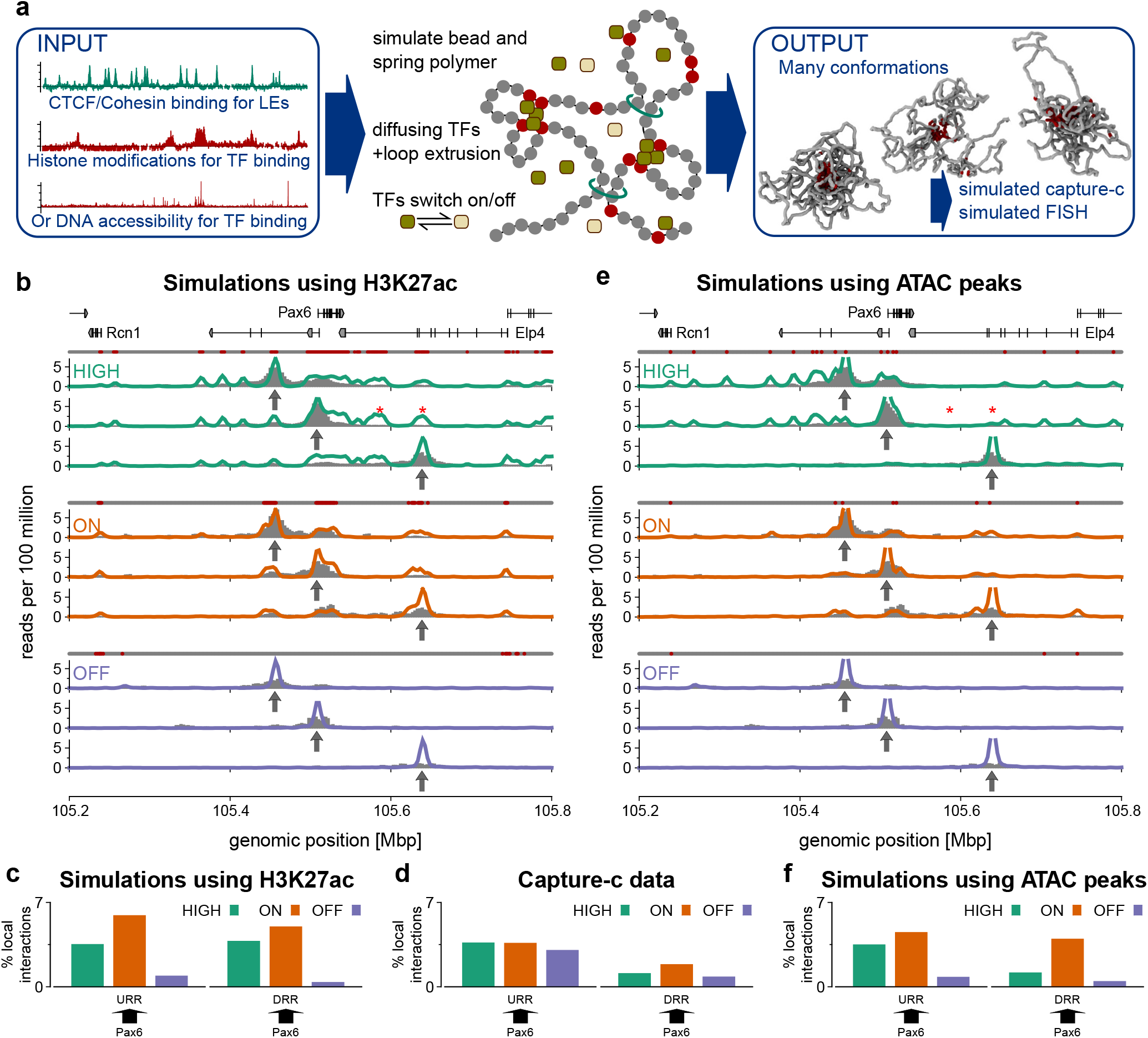
Polymer simulations predict chromosome interactions in cell lines. **(a)** Schematic of the simulation model. A bead-and-spring polymer covers a 5 Mbp region around Pax6; each bead represented 1 kbp of chromatin. Initially ChIP data for H3K27ac was used to colour (mark) beads, and CTCF/Rad21 data were used to identify loop anchor beads. Later analysis used ATAC-seq data to colour beads. Freely diffusing beads represented TFs and bind coloured polymer beads; these switched back and forward between a binding and non-binding state. LE factors (represented by additional springs in simulations, and shown as red rings) bind at adjacent polymer beads and extrude loops; extrusion was halted if the LE met an anchor bead which was orientated against the direction of extrusion. LEs were removed from (and returned to) the polymer at a constant rate. The model was used to generate a population of conformations from which simulated Capture-C and FISH measurements were obtained. Full details are given in Methods. **(b)** Simulated Capture-C profiles (coloured lines) are shown for three cell lines, for three different viewpoints (at the URR, DRR and Pax6, indicated by arrows). The corresponding experimental data is shown as grey bars. Above each set of plots a line of points show how beads were coloured as non-binding (grey) or binding (red) for TFs. In simulations Pax6 interacts strongly with a broad acetylated region downstream of the gene (red stars); this is not observed in the experimental data. **(c)** Plot showing the level of interaction between a viewpoint and a specified 10 kbp region. The arrow indicates the viewpoint (arrow base) and the interacting region (arrow head). The hight of the bar shows the number of interactions with the 10 kbp region as a percentage of interactions with the locus as a whole (chr2:105,200,000-105,800,00). **(d)** Similar plots were obtained from experimental Capture-C data. **(e)** Simulated Capture-C profiles for a model where ATAC-seq data (Fig. S3) were used to infer TF binding sites instead of H3K27ac. The erroneous interactions marked in (a) are now absent (red stars). **(f)** Similar plots to those in (c) but from the simulations using ATAC-seq data.

To validate the model we used the Capture-C protocol (a combination of 3C and oligonucleotide capture followed by high-throughput sequencing^24^) to obtain interaction profiles from a set of probes, or “viewpoints” across the locus (see Methods and Fig. S2). The simulations gave good predictions of chromatin interactions in the OFF and ON cell lines (Fig. 1); they showed that in the ON cells the *Pax6* promoters interact with both distal enhancers. However, notably, the model failed to correctly predict chromatin contacts in the HIGH activity cells (Fig. 1b).

### DNA accessibility better predict locus folding

In the Pax6-HIGH cells there was a reduction in looping to the DRR compared to the ON cells, despite a broadening of the H3K27ac mark; this is inconsistent with the typical looping model for enhancer action, and is not correctly predicted by simulations (which did show promoter-enhancer loops, as well as interactions with other acetylated regions between *Pax6* and the DRR - red stars in Fig. 1b). From this analysis a simple promoter-enhancer looping mechanism for regulating chromatin folding does not occur. Our previous studies on globin loci^5^ use DNA accessibility data as a proxy for protein binding. In addition to mapping epigenetic marks ENCODE has extensively characterised chromatin disruptions using both DNaseI sensitivity and ATAC-seq; to determine if this information gave improved model predictions we generated ATAC-seq data on the three cell lines (Fig S3). This revealed that in the ON cells there is an ATAC peak within the DRR, whereas in the Pax6-HIGH cells this peak is absent (despite the broadening of the H3K27ac mark). Simulations predicated on TFs only binding to ATAC peaks gave better predictions of Capture-C data in all three cell lines (Fig. 1e-f).

These results show that while histone modification data can be used to recover the large-scale domain structure of chromosome organisation^6,7^, for more complex loci at higher resolution, DNA accessibility gives a better prediction of TF binding driven structure. The molecular basis for this is unclear but probably reflects a more direct correlation between TFs binding and disrupted chromatin, whilst histone modifications are generated indirectly as a consequence of acetyltransferases or methylases binding to chromatin.

### High level transcription drives local decompaction

From the panoply of simulated structures individual measurements can be extracted equivalent to information obtained from FISH experiments (Fig 2). This provides values for the distance between pairs of points on the polymer but also enables the volume of the locus to be predicted. Initially compaction upstream and downstream of the *Pax6* locus was measured (using pairs of FISH probes at URR/Pax6 and Pax6/DRR respectively, Fig. 2). To validate these results we performed 3-D FISH experiments. Both probe pairs showed a non-monotonic variation in separation as a function of *Pax6* activity, both *in vivo* and in simulations. Separations were reduced in Pax6-ON compared to OFF cells, but were larger in Pax6-HIGH compared to Pax6-ON. To quantify how well the simulations predicted the data a measure called the K-score was defined (see Methods and Fig. S4), ranging from zero to one (from no agreement to perfect overlap of separation distributions); despite a high score of K=0.70 simulations did not correctly predict the result that Pax6-HIGH cells showed significantly larger probe separations than Pax6-OFF. Varying bead size in the simulations to either 400 bp or 3 kb reduced the K-score (Fig. S4).

**Figure 2.**
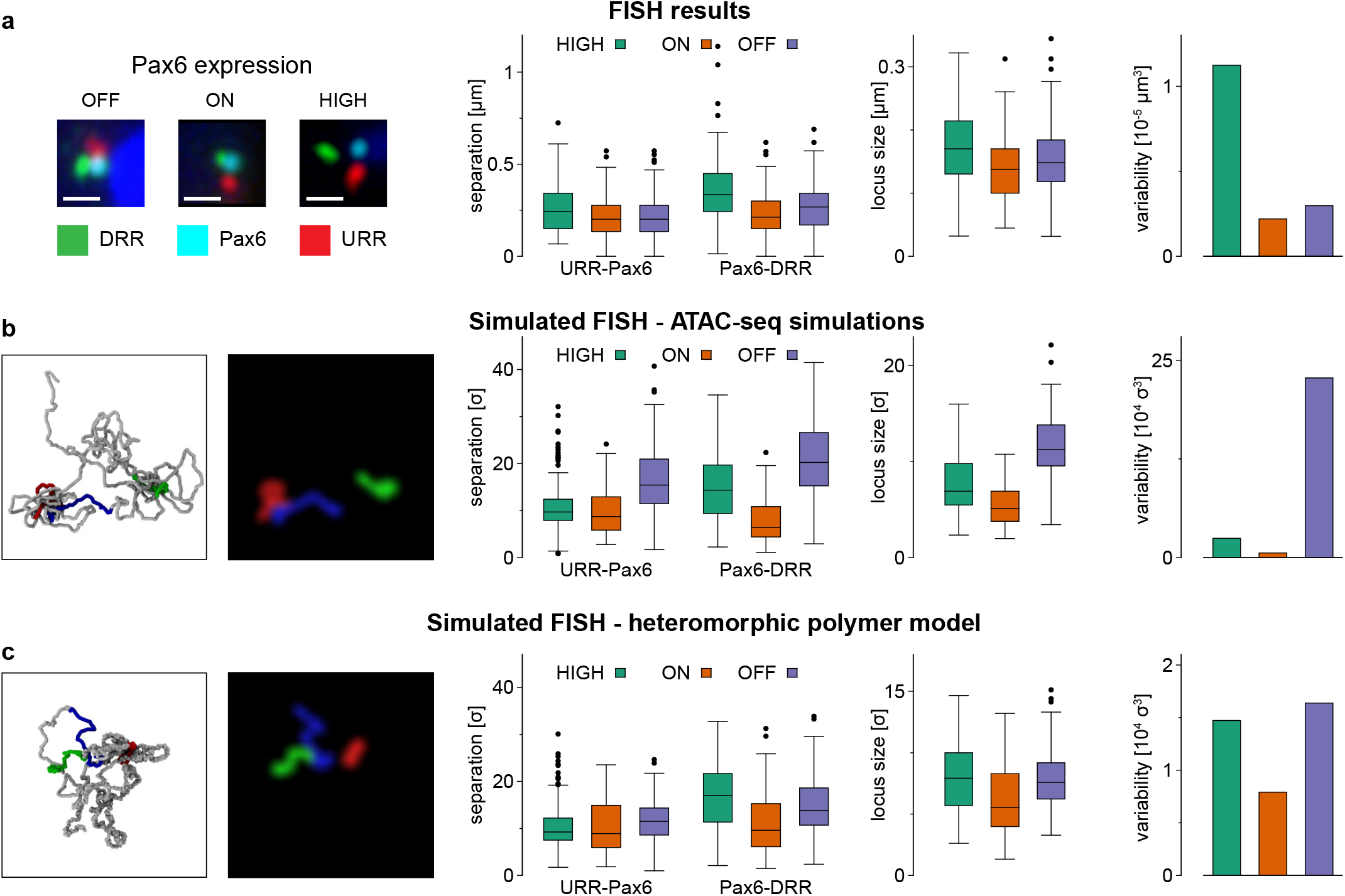
Fluorescence microscopy gives single cell information on locus conformation. **(a)** Results from 3-D FISH experiments using probes positioned at the URR, DRR and Pax6 (Fig S5a) LEFT: Representative three-colour 3-D FISH images. Scale bar is 0.5 μm. MID-LEFT: Distributions of probe separations are shown as box plots. MID-RIGHT: Three-colour experiments giving simultaneous separations of three probes allowed a measure of the size of the locus to be calculated (see Methods and Fig. S5c,d). Distributions of this measure are shown as box plots. RIGHT: We also calculated a measure of the variability of the locus conformation (see Methods and Fig. S5e-j). In general the locus becomes more compact in Pax6-ON cells compared to Pax6-OFF, but becomes less compact (and more variable) in Pax6-HIGH cells. **(b)** Similar plots were obtained from the locus conformations from the simulations using ATAC data to define TF binding sites (as shown in Fig. 1e-f). The leftmost panel shows a representative snapshot of the locus conformation, alongside a simulated FISH image shown for illustrative purposes. In box plots lengths are given in simulation units σ (see Methods). Simulation predictions depart from the experimental measurement in that here the Pax6-OFF cells are the least compact and most variable. **(c)** Simulated FISH measurements from the new heteromorphic polymer model. Agreement with the experimental data improves in that the locus is least compact in Pax6-HIGH cells, and the variability is also large in those cells.

To test how the simulations predicted overall locus volume, the volume enclosed by three FISH probes was experimentally measured (URR/*Pax6*/DRR, see Methods). Our previous studies^2^ showed that transcriptionally active regions are decompacted; surprisingly the simulations predicted that the HIGH cells would be more compact than Pax6-OFF. This was not consistent with values obtained from 3-probe FISH (Fig. 2; note also that again a non-montonic trend through OFF-ON-HIGH cells was observed). We also designed a measure of cell-to-cell variability by computing the level of the scatter in a plot where simultaneous URR-*Pax6*, Pax-DRR and URR-DRR measurements were shown on three axes (Fig. 2a, right and Fig. S5e-g; see Methods). Simulations predicted that Pax6-OFF cells would show most variability; again this was inconsistent with 3-probe FISH, where Pax6-HIGH showed most variability.

### Polymer simulations of heteromorphic chromatin fibres predict experimental data

The model described above agreed with population based 3C-style data, but could not accurately predict trends observed in single cell FISH. We reasoned that because our model assumed a homomorphic fibre, variation between cell types could only arise from differences in the TF binding or CTCF locations as loop extruder anchor sites. However, it is known that chromatin fibres can adopt alternate configurations^1,2,25,26^, and recent RICC-seq experiments suggest there are two main local structural motifs, associated with chromatin fibres: a more open and a more compact conformation^27^. It has also been suggested that acetylation marks regions of disrupted chromatin^27–29^, and indeed RICC-seq data showed a more open chromatin structure was correlated with H3K27ac^27^. Consistent with this, volumes measured for individual FISH probes were correlated with the level of H3K27ac within the probe region (Fig. S5b, i). Therefore we hypothesised that including a different chromatin fibre structure at H3K27ac regions might improve the agreement between simulations and FISH data. To achieve this in a simple way additional springs were introduced to regions which do not have the acetylation mark — i.e., where the fibre had a higher linear compaction (kbp of DNA in a given length of fibre) — leaving H3K27ac regions less compact (see schematic in Fig. 3a and Methods for details). We call this the highly predictive heteromorphic polymer (HiP-HoP) model.

**Figure 3.**
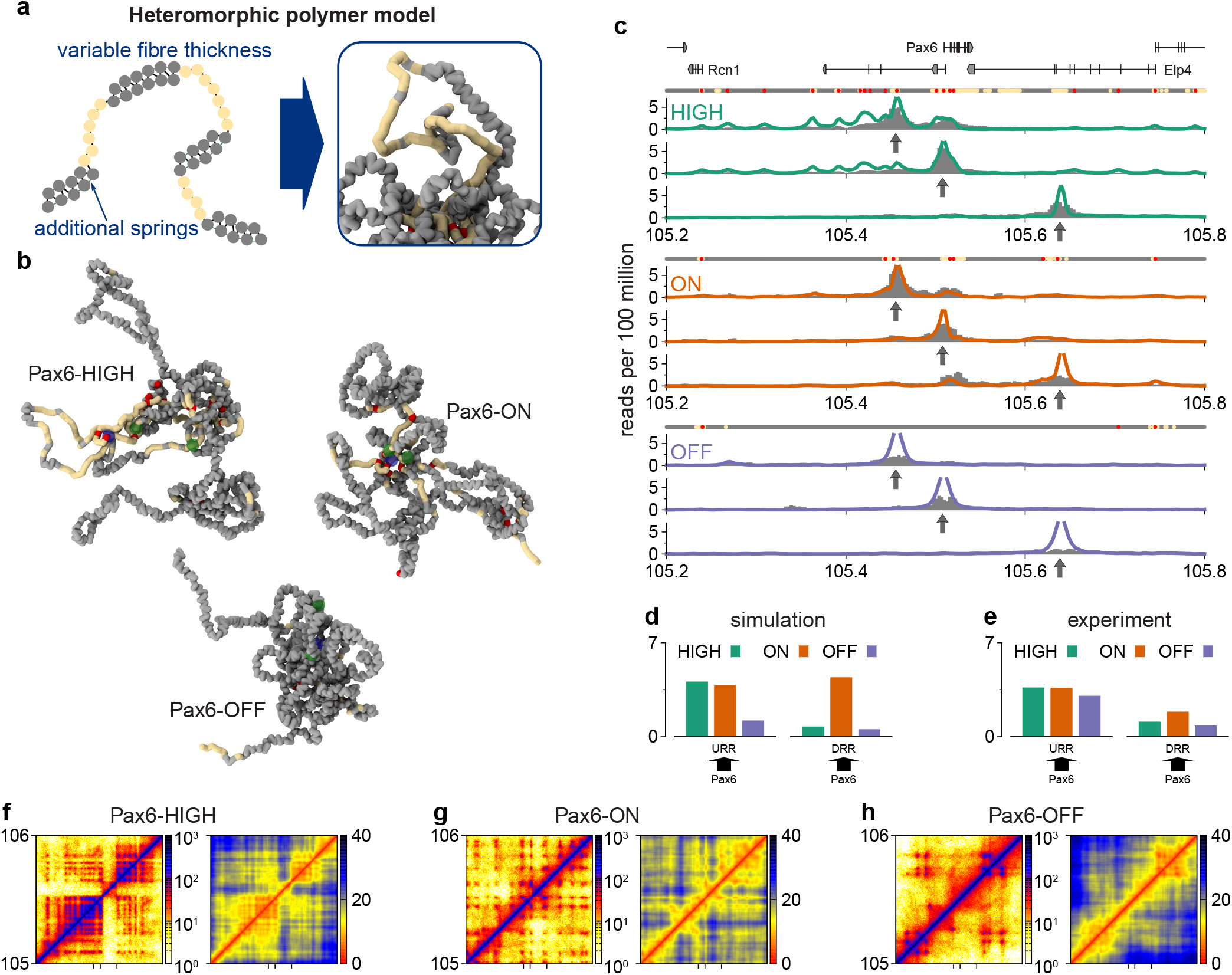
Heteromorphic chromatin fibre model gave better predictions of experimental observations. **(a)** Left: schematic showing how two levels of chromatin fibre thickness were simulated by adding additional springs between next-nearest neighbouring beads. Regions which have the H3K27ac mark (yellow) did not have these extra springs. Diffusing TFs and LEs were then added as before. Right: snapshot of a typical simulated fibre; red regions correspond to TF binding sites as inferred from ATAC peaks. TFs are not shown. **(b)** Typical snapshots of the simulated fibre in each of the three cell types. Only the region chr2:105,000,000-,106,000,000 is shown; TFs are not shown. Yellow and red regions indicate H3K27ac and ATAC regions respectively. The transparent blue sphere indicates the Pax6 promoters and green spheres indicate the URR and DRR. **(c)** Simulated Capture-C tracks from the three cell types (solid lines) for three viewpoints (positions indicated with arrows); grey bars show experimental data. Viewpoints are indicated with arrows, and the bead colours are indicated by rows of points above each set of plots: grey regions have a more compact fibre, yellow indicates the more open fibre (H3K27ac marked) and red indicates ATAC-seq peaks (TF binding). **(d-e)** Bar plots as in Fig. 1c,f showing interactions between specified viewpoints (arrow base) and a 10 kbp region around a feature of interest (arrow head). The trends seen in the experimental data (e) are now better predicted by the simulations. **(f-h)** LEFT: Simulated HiC maps. RIGHT: A similar map shows the mean distance between each bead in the region in simulation length units, sigma.

Since we do not know how chromatin structure actually varies along the fibre at high resolution, and in reality there is likely to be more than two levels, we could not expect the model to predict the FISH data exactly. Nevertheless HiP-HoP simulations did correctly reproduce all observed trends in our experiments. Most notably, we found that with this new model, the *Pax6*/DRR separations were on average furthest apart in the Pax6-HIGH cells, and this cell type also showed much higher cell-to-cell variability than in the previous simulations. The K-score (see Methods) increased to 0.77 (by nearly 10%) compared to earlier simulations (Fig. 1e), indicating improved agreement with the data (Fig S4). We defined a second quantitative metric - the Q-score - which measures the agreement between experimental and simulated Capture-C profiles. Although we noted very little visible change in the simulated Capture-C profiles, the Q-score increased by 13% to 0.51 (Fig. 1e, also see Fig S6), the extent of chromatin decompaction in Pax6-HIGH cells is apparent from inspection of simulation snapshots of the locus structure (Fig. 3b, and Video 1 and 2).

Although the Capture-C protocol only gives information about interactions for specific probed “viewpoints”, a signature of a domain structure was present in the data as we observed that probes from the left of the locus to interact more with regions towards the right, and *vice versa* (Fig. S2e). The same directional bias was found in simulated Capture-C data for Pax6-HIGH cells, but was not present in the other two cell lines (Fig. S7c). However, a full HiC like interaction map can be extracted from the simulations, and these indeed showed domains for all cell types (Fig. 3f-h and S7b). A reason for the seemingly different signal might be that the capture targets (viewpoints) tend to be positioned on TF binding beads, and over estimation by the HiP-HoP model of long range interactions at these sites may skew the directionality measure.

The simulation scheme also allows investigation of different scenarios for genome organisation. For example, simulations can be performed with different aspects of the model switched off (Fig. S8). We found that switching off LEs (similar to a knock-down of cohesin or its loader, leaving only diffusing TFs binding to ATAC peaks and the heteromorphic polymer based on H3K27ac data) led to a reduction in agreement with data (the Q-score reduced by 20%, FigS6; and the FISH K-score by 9%, Fig S4). At a larger scale, the simulated HiC maps changed: they became more “spotty” (showing promoter-enhancer interactions) and the domains less prominent (Fig. S8b). If instead TFs were removed from the model (keeping LEs and the heteromorphic polymer) the domains remain, but most enhancer-promoter interactions disappear (Fig. S8c); also the Q-score reduces by about 15% compared to the simulations in Fig. 3, although the K-score shows a small increase of about 2%). This points to a scenario where TFs give rise to promoter-enhancer interactions, while LEs generate domains.

### HiP-HoP simulations reveal multiple 3-D chromatin structures for the *Pax6* locus

Experimental and simulated FISH measurements suggest that there is substantial variation in the distribution of inter-probe distances in Pax6-HIGH cells. More detailed information on structural variability is best extracted from an analysis of individual simulated structures, as these give information about chromatin fibre conformation at high resolution (single monomer, or 1 kbp). We focus here on the case of Pax6-HIGH cells. First, a qualitative inspection of simulated conformations (see Video 3-5) provided a striking visual impression of the large structural diversity in *Pax6* folding - it was apparent that the shape and size of the locus varies widely. Second, a clustering analysis of all simulated structures based on mean-squared differences of monomer separations (see Methods) showed that there are multiple typical structures for the chromatin fibre around *Pax6* (Fig. 4), with several possible structure classes (Fig. 4a). The distribution of locus size and shape (quantified via the radius of gyration and shape anisotropy) showed only slight variation between each class (Fig. 4b-c; the largest difference is between class A, where the locus is usually larger and more elongated, and class E, where the locus is smaller and more spherical). Intriguingly, each class contained some structures where *Pax6* and its distal enhancers are in contact, and some where they are far apart (Fig. 4d). These two motifs are likely to be associated with different levels of transcriptional activity, yet our analysis shows that this is largely independent of the larger-scale structure of the locus. While a similar clustering analysis has pointed to structural diversity in the folding of other chromatin loci, the extent of variability found in the *Pax6* locus exceeds that observed in less complex loci such as globin (where simulations suggested the existence of two main classes of structures^5^).

**Figure 4.**
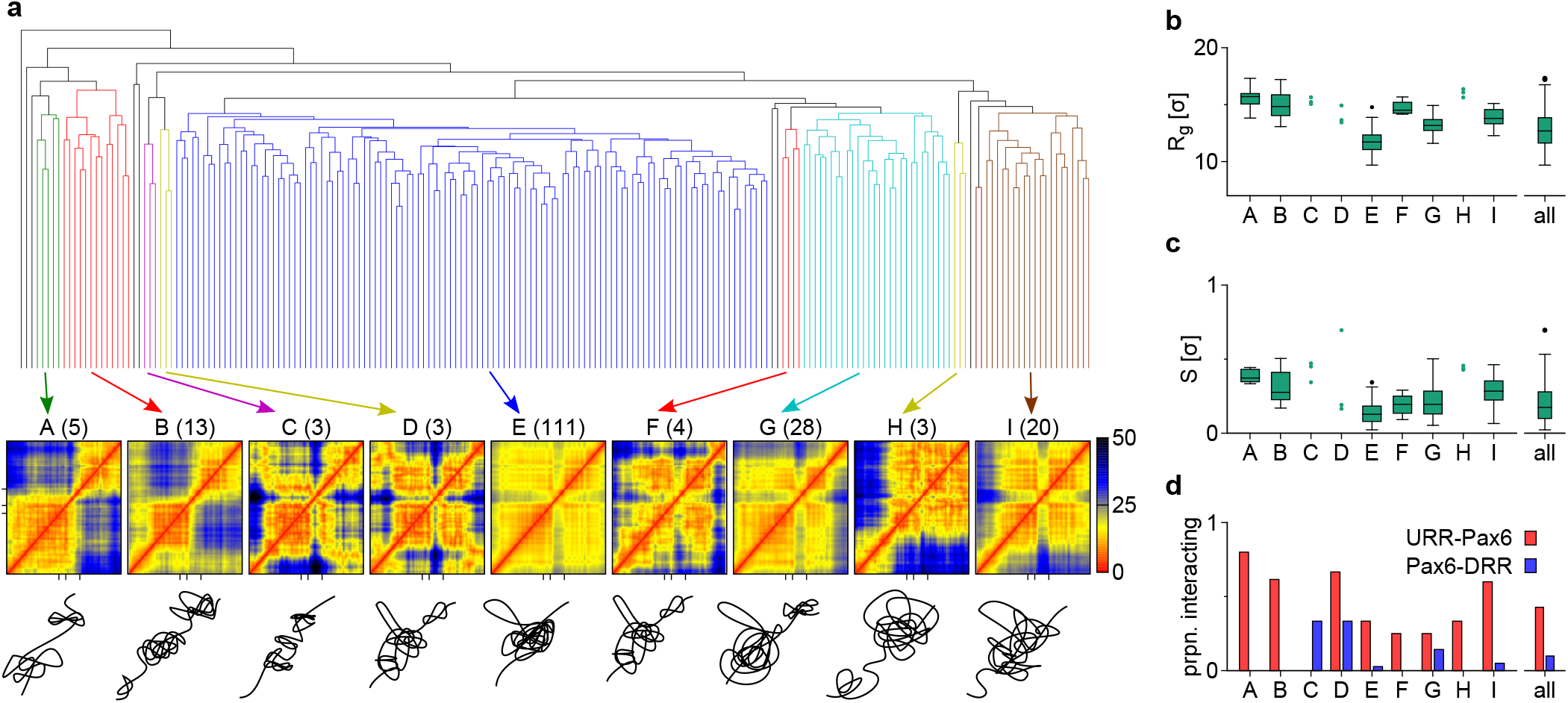
Hierarchical clustering analysis of 200 simulated Pax6-HIGH conformations revealed groups of similar structures. **(a)** TOP: Dendrogram generated via hierarchical clustering using an ‘average’ linking criterion. A Euclidean distance metric based on the pairwise difference between the separations of all pairs of beads was used (see Methods). Some groups of similar conformations are highlighted in colour. MIDDLE: For each highlighted group a distance map of the region chr2:105,000,000-106,000,000 is shown (colour scale gives the distance between pairs of beads in simulation distance units σ). Axis ticks on the bottom of the plots show the positions of the URR, Pax6 promoters, and the DRR. BOTTOM: Sketched representations of potential combinations. **(b)** The distribution of the radius of gyration of the region chr2:105,000,000-106,000,000 is shown as a box plot for groups highlighted in (a). This is a measure of the size of the locus. Where there are fewer than four conformations in a group single points are shown. To the left the distribution for all 200 conformations is shown. **(c)** Similar plot to (b) but showing the shape anisotropy, a measure of the relative shape of the locus. This ranges from 0, for a spherically symmetric arrangement, to 1 for a linear arrangement. **(d)** For each group of conformations highlighted in (a) the proportion in which Pax6 interacts with each of the distal regulatory regions is indicated. As expected from the simulated Capture-C data shown in Fig. 3, in Pax6-HIGH cells interactions with the DRR are rare.

Another way to group the simulated structures is to directly consider the interactions between the distal regulatory regions and the *Pax6* promoters (Fig. S9). There were five possible combinations: none of the elements were in contact, *Pax6* contacted one of the enhancer regions, the two enhancer regions were in contact with each other but not *Pax6*, or all three were in contact. Interestingly, within the 200 structures for each cell type, none had interactions between the two enhancers without them also interacting with *Pax6*, and the DRR never interacted with *Pax6* in isolation. Consistent with the observations detailed above, in Pax6-OFF cells the majority of conformations had no Pax6-enhancer interactions, in Pax6-ON cells a large proportion of conformations had *Pax6* interacting with both enhancers, whereas in Pax6-HIGH cells *Pax6* more often only interacted with the URR (Fig. S9a). Consistent with the clustering analysis, the *Pax6* interactions did not depend on the size or shape of the locus as a whole (Fig. S9b-c).

### Application of HiP-HoP simulations to active chromatin loci

Above we focussed on *Pax6* folding, but since our predictive heteromorphic polymer simulation approach only requires four different datasets as input (DNA accessibility, histone acetylation and CTCF/Rad21), it is applicable to a large number of active chromatin loci in different cell lines (associated with different levels of locus activity). To test how well the HiP-HoP model performs at other loci, we studied the folding of the α and β-globin loci in mouse erythroid cells, which involve simpler genomic interactions with respect to *Pax6^5^*. As expected, our simulated structures for globin compared favourably with previously published Capture-C and FISH data (Fig. 5 and S10).

To show that the HiP-HoP model can work in different organisms, we also considered the human Sox2 locus, a key reprogramming gene (Fig. S11). We found good agreement with experimental HiC contact maps in stem cells and umbilical vein epithelial cells (HUVECs). These results show that our model is portable to other loci, and that it can be used to predict folding of loci which have not yet been investigated either by chromosome conformation capture or FISH.

**Figure 5:**
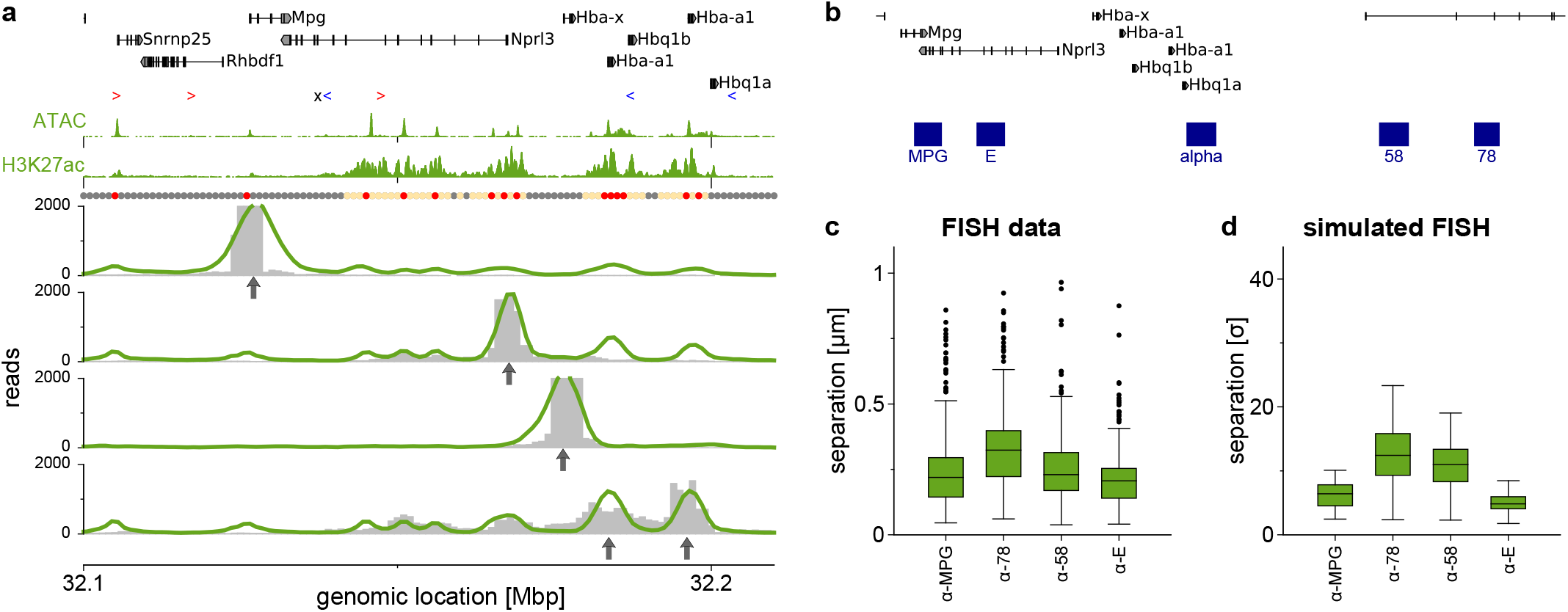
The α globin locus. Results from simulations of a 2~Mbp region around the α globin gene in mouse erythroid cells using the HiP-HoP model. **(a)** Capture-C data from four viewpoints (indicated with arrows). Simulation data is shown as a solid line, whereas experimental data (from ref 24) is shown as grey bars. Since there are two copies of the α globin gene, oligo captured fragments could have come from either copy, hence two viewpoints are shown on the same plot. A map of the locus, the data used as simulation input, and points indicating the bead colours are shown above. **(b)** Map of the locus showing positions of the FISH probes. **(c)** FISH data (obtained from ref 5). **(d)** Simulated FISH data using the same probe pairs.

## Discussion

In this study we developed a new simulation model of 3-D genome organisation modelling chromatin as a heteromorphic polymer, where the H3K27ac histone modification is associated with a locally disrupted chromatin fibre; this was corroborated by a strong positive correlation between histone acetylation and putative regions of disrupted chromatin based on RICC-seq experiments^27,29^. The failure of simple models and the development of novel simulations led to greater understanding of the *Pax6* locus; a substantial increase in *Pax6* expression observed in the HIGH activity cells is *not* accompanied by an increase in looping interactions between the *Pax6* promoters and the DRR (a change which is observed when going from Pax6-OFF to Pax6-ON cells). Additionally, in Pax6-HIGH cells microscopy and HiP-HoP simulations showed that the chromatin fibre at the DRR must undergo a dramatic decondensation, associated with a 50% increase in the mean separation between this enhancer site and the *Pax6* promoter. Two possible explanations for these results are that either the downstream enhancer is not involved in up-regulation of *Pax6* in these cells, or that there is an enhancer activity which does not require physical proximity to the promoter, but is instead associated with chromatin decompaction^30^. One can speculate on possible mechanisms through which decompaction of an enhancer region might lead to up-regulation of a nearby gene: perhaps the region adopts a fibre structure which can more readily accommodate supercoiling generated by a transcribing polymerase; alternatively transcription at the enhancer itself might lead to a localization of activating proteins which facilitates transcription at the promoter; or perhaps the expansion of the enhancer region alters the dynamical properties of the wider locus. Whatever the mechanism, an interesting feature at *Pax6* is that the DRR seems to operate differently in different cell lines, an area for future research.

These simulations also suggested that for *Pax6* (and possibly other complex loci) there is a large degree of cell-to-cell variation in locus conformation. This is different to the case of the globin loci where both loci were found to adopt one of a small number of preferred configurations^5^. Earlier computational models failed to predict trends in the FISH data, but gave similar predictions for Capture-C profiles; this highlights a potential issue for approaches that generate conformations based on fitting to existing HiC data - more than one solution may be possible.

In the future it would be informative to apply HiP-HoP to many different loci in different cell types to understand the organisational principles of different classes of gene. We also expect the model to be readily extended to account for co-localisation of repressed regions. As input to HiP-HoP is based on widely available datasets — ATAC or DNAase-seq for TF binding, H3K27ac to predict disrupted chromatin regions, and ChIP data for CTCF and the cohesin subunit Rad21 to define loop anchors —the same model will be applicable to active loci generally. Indeed, we have shown that these predictive heteromorphic polymer simulations can be successful applied to loci in different cell types and organisms, such as α and β-globin, *Sox2*, in stem cells and tissue derived cell lines, and in human or mouse. Unlike other approaches that require 3C based data as an input, the HiP-HoP model does not require this data making it well suited for predicting chromatin structure at promoters and enhancers genome-wide.

## Acknowledgements

We thank members of the Gilbert and Marenduzzo labs for helpful discussions and Dirk-Jan Kleinjan for support in the early stages of the project. N.G. is an MRC Senior Resarch Fellow (MR/J00913X/1) and research in the D.M. lab is supported by ERC (CoG 648050, THREEDCELLPHYSICS).

## Author contributions

A.B., S.B. undertook experiments, C.B. carried out simulations. N.G. and D.M. conceived the project and all authors contributed to writing the manuscript

## Competing Financial Interests

The authors declare no competing financial interests

